# Dysregulation of Kappa Opioid Receptor Neuromodulation of Lateral Habenula Synaptic Function following a Repetitive Mild Traumatic Brain Injury

**DOI:** 10.1101/2024.05.01.592017

**Authors:** William J. Flerlage, Sarah C. Simmons, Emily H. Thomas, Shawn Gouty, Brian M. Cox, Fereshteh S. Nugent

## Abstract

Mild traumatic brain injury (mTBI) increases the risk of cognitive deficits, affective disorders, anxiety and substance use disorder in affected individuals. Substantial evidence suggests a critical role for the lateral habenula (LHb) in pathophysiology of psychiatric disorders. Recently, we demonstrated a causal link between persistent mTBI-induced LHb hyperactivity due to synaptic excitation/inhibition (E/I) imbalance and motivational deficits in self-care grooming behavior in young adult male mice using a repetitive closed head injury mTBI model. One of the major neuromodulatory systems that is responsive to traumatic brain and spinal cord injuries, influences affective states and also modulates LHb activity is the dynorphin/kappa opioid receptor (Dyn/KOR) system. However, the effects of mTBI on KOR neuromodulation of LHb function is unknown. To address this, we first used retrograde tracing to anatomically verify that the mouse LHb indeed receives Dyn/KOR expressing projections. We identified several major KOR-expressing and Dyn-expressing synaptic inputs projecting to the mouse LHb. We then functionally evaluated the effects of *in vitro* KOR modulation of spontaneous synaptic activity within the LHb of male and female sham and mTBI mice at 4week post-injury using the repetitive closed head injury mTBI model. Similar to what we previously reported in the LHb of male mTBI mice, mTBI presynaptically diminished spontaneous synaptic activity onto LHb neurons, while shifting synaptic E/I toward excitation in female mouse LHb. Furthermore, KOR activation in either mouse male/female LHb generally suppressed spontaneous glutamatergic transmission without altering GABAergic transmission, resulting in a significant reduction in E/I ratios and decreased excitatory synaptic drive to LHb neurons of male and female sham mice. Interestingly following mTBI, while responses to KOR activation at LHb glutamatergic synapses were observed comparable to those of sham, LHb GABAergic synapses acquired an additional sensitivity to KOR-mediated inhibition. Thus, in contrast to sham LHb, we observed a reduction in GABA release probability in response to KOR stimulation in mTBI LHb, resulting in a chronic loss of KOR-mediated net synaptic inhibition within the LHb. Overall, our findings uncovered the previously unknown sources of major Dyn/KOR-expressing synaptic inputs projecting to the mouse LHb. Further, we demonstrate that an engagement of intra-LHb Dyn/KOR signaling provides a global suppression of excitatory synaptic drive to the mouse LHb which could act as an inhibitory braking mechanism to prevent LHb hyperexcitability. The additional engagement of KOR-mediated modulatory action on LHb GABAergic transmission by mTBI could contribute to the E/I imbalance after mTBI, with Dyn/KOR signaling serving as a disinhibitory mechanism for LHb neurons in male and female mTBI mice.

## 1. Introduction

Mild traumatic brain injury (mTBI) is a significant public health burden due to the high risk of long-term detrimental effects of the injury including cognitive deficits, affective disorders such as depression, anxiety and substance use disorders^1, 2^. A major neurmodulatory stress system that could play an important role in mood dysregulation associated with mTBI is dynorphin (an endogenous opioid ligand) and its cognate receptor, kappa opioid receptor (KOR) as this system may be engaged following traumatic brain and spinal cord injuries ^3–16^. However, brain region-specific modulation of Dyn/KOR system following mTBI has yet to be explored. Dyn/KOR systems are widely expressed in stress– and reward/ motivational brain circuits, regulates motivational states and behavioral responses to emotional and stressful situations and their dysregulation is implicated in the development of mood disorders, including depression and anxiety, as well as in drug addiction ^17–20^. Here, we investigated a possible mTBI-induced dysregulation of Dyn/KOR-mediated signaling within the lateral habenula (LHb), a major node for mood and motivation regulation that is also responsive to Dyn/KOR system.

Several lines of evidence from clinical and preclinical studies indicate that the LHb is critically involved in emotional and mood regulation, reward-based decision making, avoidance of stressful situations/stimuli, and stress responses. Compelling evidence indicates that LHb dysfunction contributes to long-term mood, motivation and emotional dysregulation^21–30^. LHb neurons respond to omission of expected rewards and unpleasant/aversive/stressful events, thereby encoding reward omission, aversion and punishment signals and promoting avoidance behaviors through suppression of monoaminergic systems involved in emotion, reward, motivation, and cognitive processing. LHb receives diverse inputs from forebrain limbic and basal ganglia regions such as medial prefrontal cortex (mPFC), entopeduncular nucleus (EP), ventral pallidum (VP), bed nucleus of stria terminalis (BNST as a part of extended amygdala), ventral tegmental area (VTA), periaqueductal gray (PAG), lateral preoptic area (LPO) and lateral hypothalamus (LHA) and projects to different midbrain structures including rostromedial tegmental area (RMTg), VTA, dorsal raphe nucleus (DRN), PAG and locus coeruleus (LC). The majority of glutamatergic outputs of LHb synapses on GABAergic interneurons of the VTA and DRN as well as RMTg GABAergic neurons projecting to VTA and DRN, thus LHb glutamatergic activity indirectly provides a potent inhibitory drive suppressing VTA dopaminergic and raphe serotonergic neuronal activities. Hyperactivity of LHb neurons is generally presumed to serve as a critical neural substrate for depression and other psychiatric illnesses, although LHb hypofunction/loss of proper LHb functioning can also mediate negative affective states, social deficits, risky decision-making and impulsivity^29–35^.

While there is substantial evidence for direct Dyn/KOR regulation of the mesocorticolimbic system^36^, less is known about the role of Dyn/KOR-mediated regulation of LHb in normal LHb function or its dysregulation within the LHb under pathological conditions. Previous work from our laboratory provided the first demonstration of direct Dyn/KOR-mediated modulation of LHb neuronal excitability and synaptic function in male rats in which we also showed how a severe early life stress (maternal deprivation) in male rats alters LHb sensitivity to this system^37^. Our study suggested a functional Dyn/KOR system within the LHb, though it was unclear whether KOR-mediated synaptic modulation also exists in the mouse LHb and is responsive to mTBI injuries. Interestingly, a recent study showed that a repeated blast injury mTBI model in male mice increases dynorphin A levels in the mouse brain and that systemic pretreatment of male mice with a long-acting selective KOR antagonist (nor-BNI) blocks long-term aversive and anxiety-like behaviors in blast-injured male mice one month post-injury^38^. This raises the possibility that mTBI engagement of Dyn/KOR system may contribute to the development of persistent mood-related behavioral deficits following this model of mTBI. However, it is yet to be identified which Dyn– and KOR-expressing neurons within brain reward/stress circuits are engaged and responsive to mTBI injuries. Our recent work has also highlighted a causal role between LHb hyperactivity and behavioral deficits in self-care grooming behavior in male mice following a repetitive closed head injury mTBI model^39^. We found that a repetitive closed head injury model of mTBI increases LHb spontaneous activity through a shift in synaptic excitation to inhibition (E/I) balance towards excitation in parallel with deficits in self-care grooming, while chemogenetic silencing of LHb neurons reverses motivational deficits in grooming behaviors in male mice. Furthermore, we also found that mTBI similarly increased LHb neuronal excitability in both male and female mice, partly through increased corticotropin releasing factor (CRF)/CRF receptor 1 signaling within the LHb^40^. Here, we used viral-based retrograde tracing to first anatomically identify Dyn– and KOR-expressing projections to the LHb and then functionally examined the impact of mTBI on KOR-modulation of spontaneous glutamatergic and GABAergic transmission to LHb neurons in male and female mice. Our findings uncovered several major brain regions that can provide Dyn/KOR signaling to the LHb. Furthermore, we provide evidence for a functional KOR-mediated suppression of excitatory synaptic drive to the LHb that could act as an inhibitory neuromodulatory braking mechanism to inhibit LHb activity or prevent LHb hyperactivity. We show that additional engagement of KOR-mediated modulatory action on LHb GABAergic transmission could lead to pathological imbalance of excitation and inhibition as we observe in male and female mouse LHb following mTBI. Overall, our study provides evidence for protective role of Dyn/KOR-mediated regulation of LHb activity that is susceptible to mTBI injury and may contribute to E/I imbalance and LHb hyperactivity in mice that sustained injury.

## 2. Materials and Methods

### 2.1. Animals

All experiments were carried out in accordance with the National Institutes of Health (NIH) *Guide for the Care and Use of Laboratory Animals* and were approved by the Uniformed Services University Institutional Animal Care and Use Committee. C57BL/6 male mice, Pdyn (prodynorphin) Cre mice (B6;129S-*Pdyn^tm^*^1.1^(cre)*^Mjkr^*/LowlJ, Strain #:027958) and KOR Cre (*Oprk1^tm^*^1.1^(cre)*^Sros^*/J, Strain #:035045) mice (Jackson Laboratory) were acquired at ∼postnatal day 35-49 (PN35-P49) and allowed at least 72hrs of acclimation before the initiation of experimental procedures. Mice were group-housed in standard cages under a 12hr/12hr light-dark cycle with standard laboratory lighting conditions (lights on, 0600-1800, ∼200lux), with ad libitum access to food and water. All procedures were performed beginning 2– 4hr after the start of the light-cycle. All efforts were made to minimize animal suffering and reduce the number of animals used throughout this study.

### 2.2. Virus injections for anatomical retrograde tracing

Male and female Pdyn Cre and KOR Cre mice (∼P42-50) were anesthetized with isoflurane using ∼ 1–3% vaporized isoflurane with oxygen and fixed into a stereotaxic frame. Body temperature was maintained at 37⸰C throughout the procedure and during recovery with a heating pad. The scalp was shaved and a midline incision was made on the skin overlying the skull. The retrograde adeno-associated virus (AAV) tracer (AAVrg pAAV-Ef1a-DIO EYFP, Addgene#27056) was infused (∼50 nL/side; over 5min using a Nanoject III Injector, Drummond) using pulled glass pipettes into the LHb (coordinates from bregma: AP, −1.6 mm; ML, ± 0.5 mm; DV, −3.0 mm). The injection pipette was not removed until 5min after the end of the infusion to allow diffusion of the virus. During the recovery (approx. 30-60 min), each mouse was placed in a cage that is kept warm by a heating pad (placed under the cage) or under the heating lamp to prevent hypothermia. The investigators were responsible for post-operative care, which included continuous monitoring of the mice until they were able to make purposeful movements. The health and weight of animals were monitored and stored in a medical record for a minimum of 3 days following surgery. Mice were injected subcutaneously with 5 mg/kg carprofen (for pain relief) immediately after surgery, and monitored daily for signs of pain, and carprofen was administered as needed.

### 2.3. Histochemistry and microscopy

At a minimum of three-weeks following viral injection, mice were anesthetized with an intraperitoneal injection containing ketamine (80 mg/kg) and xylazine (10 mg/kg) and perfused through the aorta with 100 ml heparinized 1x phosphate buffered saline (PBS) followed by 150 ml of 4% paraformaldehyde. The brains were dissected and placed in 4% PFA for 24hr and then cryoprotected by submersion in 20% sucrose for 3 days, frozen on dry ice and stored at –70°C until sectioned. Brain sections were cut at 20 μm, generating a series of sections ranging from Bregma AP 2.10 to –4.26. Slide mounted sections were cover-slipped using Prolong Gold with DAPI (Invitrogen: Eugene, Oregon). All sections were scanned using a Zeiss AxioScan at 10X magnification. Labeled neurons were counted across regions of interest (ROI) identified using the Allen Mouse Brain Atlas. For each ROI the sum of labeled cells from three consecutive sections were counted and an average for each ROI was calculated between subjects. Data is presented relative gradient of labeling: – none; (+) sparse <5 eYFP-labeled cells; + medium 5-25 eYFP-labeled cells; high 25-50 eYFP-labeled cells; very high >50 eYFP-labeled cells across all sections. For quantification of Dyn+− or KOR+− neurons, three mice/genotype were analyzed.

### 2.4. Repetitive mild traumatic brain injury model

Mice underwent either repeated sham or repeated closed head injury (CHI) at ∼ PN56, delivered by the Impact One, Controlled Cortical Impact (CCI) Device (Leica; Wetzler, Germany) utilizing parameters which were previously described^39^. Briefly, mice were anesthetized with isoflurane (3.5% induction/2% maintenance) and placed into a stereotaxic frame for each head impact injury. Repeated CHI-CCI in mTBI group consists of 5 discrete concussive impacts to the head delivered at 24hr intervals generated by an electromagnetically driven piston (4.0m/s velocity, 3mm impact tip diameter, a beveled flat tip, 1.0 mm depth; 200 ms dwell time) targeted to bregma as visualized through the skin overlying the skull following depilation. Sham mice underwent identical procedures except without delivery of impact. Body temperature of mice was maintained at 37^⸰^C throughout by a warming pad and isoflurane exposure and surgery duration was limited to no more than 5 minutes for each impact. Following sham or CHI-CCI surgery completion, mice were immediately placed in a supine position in a clean cage on a warming pad and the latency to self-right (righting reflex) was recorded.

### 2.5. Slice preparation

Mice were deeply anesthetized with isoflurane and immediately transcardially perfused with ice-cold artificial cerebrospinal fluid (aCSF) containing (in mM): 126 NaCl, 21.4 NaHCO_3_, 2.5 KCl, 1.2 NaH_2_PO_4_, 2.4 CaCl_2_, 1.00 MgSO_4_, 11.1 glucose, 0.4 ascorbic acid; saturated with 95% O_2_-5% CO_2_. Brain tissues were maintained on ice-cold aCSF and tissue sections containing LHb were sectioned at 220μm using a vibratome (Leica; Wetzler, Germany) and subsequently incubated in aCSF at 34 °C for at least 1– hr prior to electrophysiological experiments. For patch clamp recordings, slices were then transferred to a recording chamber and perfused with ascorbic-acid free aCSF at 28-30 °C.

### 2.6. Electrophysiology

Voltage-clamp whole-cell recordings were performed from LHb neurons in sagittal slices containing LHb using patch pipettes (3-6 MOhms) and a patch amplifier (MultiClamp 700B) under infrared-differential interference contrast microscopy. Data acquisition and analysis were carried out using DigiData 1440A, pCLAMP 10 (Molecular Devices), Clampfit, and Mini Analysis 6.0.3 (Synaptosoft, Inc.). Signals were filtered at 3 kHz and digitized at 10 kHz.

As previously described^39^, spontaneous excitatory and inhibitory postsynaptic currents (sEPSCs and sIPSCs) were recorded within the same LHb neuron in voltage clamp mode with a cesium-methanesulfonate (CsMeS)-based internal solution in intact synaptic transmission over 10 sweeps, each lasting 50 s (a total of 500s continuous recording for either sEPSC or sIPSC). Patch pipettes were filled with Cs-MeS internal solution (140 mM CsMeS, 5 mM KCl, 2 mM MgCl_2_, 2 mM ATP-Na+, 0.2 mM GTP-Na+, 0.2 mM EGTA, and 10 mM HEPES, pH 7.28, osmolality 275-280 mOsm). Cells were voltage-clamped at −55 mV to record spontaneous excitatory postsynaptic currents (sEPSCs) and +10 mV to record spontaneous inhibitory postsynaptic currents (sIPSCs) within the same neuron. The mean excitation/inhibition (sE/I) of spontaneous synaptic activity were calculated as sEPSC/sIPSC amplitude or frequency ratios from the same recording. The mean synaptic drive ratio was calculated as (sEPSC frequency × amplitude)/(sIPSC frequency × amplitude). To create cumulative probability plots for sE/I amplitude and frequency ratios and synaptic drive ratio, we randomly selected 10 sEPSC and 10 sIPSC recordings (each lasting 15s) over each sweep of the 50s recording in each cell and calculated the ratios between sEPSC and sIPSC over 15s recordings as previously described^39^. Therefore, for each cell, the combination of 10 sEPSC and 10 sIPSC yielded 100 data points (100 sE/I amplitude or frequency or synaptic drive ratios as calculated for the mean ratio values). The cell input resistance and series resistance were monitored through all the experiments and if these values changed by more than 10%, data were not included.

### 2.7. Drugs

For all experiments to activate KORs using the KOR agonist, U50,488 (Tocris, 10 µM), a within-subjects experimental design was used. Baseline recordings of sEPSCs and sIPSCs were first performed in each neuron and then the KOR agonist was added to the slice by the perfusate and sEPSCs and sIPSCs were recorded following 30-45min of drug bath application.

### 2.8. Statistics

Values are presented as mean ± SEM. The threshold for significance was set at *p < 0.05 for all analyses. All statistical analyses of data were performed using GraphPad Prism 10. For the effects of mTBI and the KOR agonist on mean values of sEPSC, sIPSC, sE/I ratios and synaptic drive ratios, two-tailed unpaired or paired Student’s t test were used, respectively. Mini Analysis software was used to detect and measure sEPSC and sIPSC amplitude and frequency (inter-event interval) using preset detection parameters of spontaneous with an amplitude cutoff of 5 pA. Effects of mTBI and the KOR agonist on the cumulative probabilities data sets were analyzed using Kolmogorov-Smirnov tests.

## 3. Results

### LHb receives Dyn– and KOR-expressing synaptic inputs from cortical and subcortical regions

Previous work from our lab demonstrated that LHb neurons are responsive to Dyn/KOR system in male rats^37^. Here we first performed retrograde anatomical tracing in transgenic male and female mice to identify LHb-projecting synaptic inputs that may provide Dyn projections to the LHb or express KORs on their synaptic terminals projecting to the LHb. We injected a retrograde-AAV tracer in the LHb (AAVrg pAAV-Ef1a-DIO EYFP, Addgene# 27056-AAVrg) in Pdyn or KOR Cre mice for Cre-dependent retrograde tracing of Dyn-/KOR-expressing neuronal projections to the LHb (Figure 1). We found only two main brain regions as the sources of Dyn projections to the LHb; subpopulations of Dyn-expressing neurons originating from mPFC or ventromedial nucleus of the hypothalamus (VMH). On the other hand, LHb-projecting KOR expressing neurons were abundant, originating from diverse forebrain, limbic and midbrain regions that are known to project to the LHb including mPFC, ventral pallidum (VP), bed nucleus of stria terminalis (BNST), entopeduncular nucleus (EP/ globus pallidus internus, GPi), extended amygdala, paraventricular thalamus (PVT), hypothalamic nuclei, zona incerta (ZI) and midbrain structures including PAG, midline nuclei (the rostral linear nucleus of the raphe, RLi and interfasicular nucleus, IF) and VTA.

**Figure 1.**
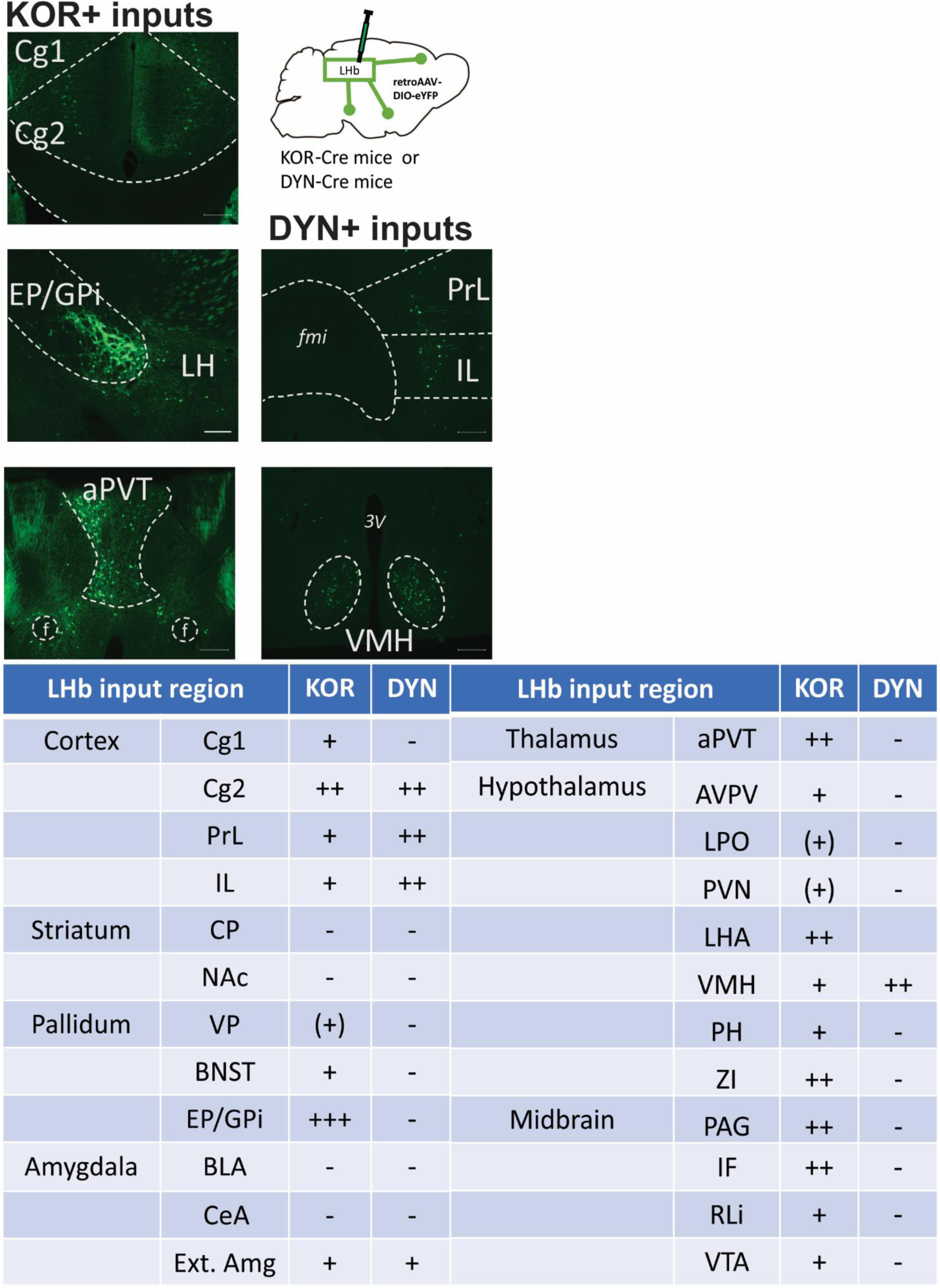
Example labeled regions from KOR-Cre (left column) and Dyn-Cre mice with LHb injection with retro-AAV-DIO-eYFP. Scale bar: 200μm. Table displays relative gradient of labeling: – none; (+) sparse <5 eYFP-labeled cells; + medium 5-25 eYFP-labeled cells; high 25-50 eYFP-labeled cells; very high >50 eYFP-labeled cells within the sections through the named structure across Cortex, Striatum, Pallidum, Amygdala, Thalamus, Hypothalamus and Midbrain. Cg: cingulate cortex, PrL: prelimbic, IL: infralimbic, CP: caudate putamen, NAc: nucleus accumbens, VP: ventral pallidum, BNST: bed nucleus of stria terminalis, EP/GPi (entopeduncular nucleus/globus pallidus internus), BLA: basolateral amygdala. CeA: central amygdala, Ext. Amg: extended amygdala, aPVT: anterior paraventricular thalamus, AVPV: Anteroventral periventricular nucleus, LPO: lateral preoptic area, PVN: paraventricular nucleus of hypothalamus, LHA: lateral hypothalamus, VMH: ventromedial nucleus of the hypothalamus, PH: posterior hypothalamus, ZI: zona incerta, PAG: periaqueductal gray, RLi: rostral linear nucleus of the raphe, and IF: interfasicular nucleus, VTA: ventral tegmental area

### 3.1. mTBI increases spontaneous excitatory synaptic drive onto LHb neurons in female mice

Previously, we examined the effects of mTBI on spontaneous synaptic transmission in LHb slices from male sham and mTBI adult mice at ∼3-4 weeks post-injury where we found that while mTBI induces a suppression of both spontaneous glutamatergic and GABAergic transmission onto LHb neurons, it results in a significant shift in spontaneous E/I balance and synaptic drive towards excitation^39^. Given sex differences exist in susceptibility to neurobehavioral effects of mTBI^41^, here we also tested the effects of mTBI in female mouse LHb. We recorded both spontaneous excitatory and inhibitory postsynaptic currents (sEPSCs and sIPSCs) within the same LHb neurons while holding in the voltage clamp mode at −55 mV and +10 mV, respectively. We found that although the average amplitude and frequency of sEPSCs and sIPSCs were not significantly different between female sham and mTBI mice, the cumulative probability amplitude and frequency (inter-event interval) plots of sEPSCs and sIPSCs were all significantly shifted to the right in LHb neurons of female mTBI mice compared to female sham mice (Figure 2 A-D, Kolmogorov–Smirnov tests, *p<0.05, **p<0.01, ****p<0.0001). The rightward shift in cumulative probabilities of both sEPSC and sIPSC inter-event intervals indicates an overall decrease in presynaptic release of both glutamate and GABA release onto LHb neurons; an observation we previously reported for the effects of mTBI in male mouse LHb^39^. We also observed rightward shifts in cumulative probabilities of sEPSC and sIPSC amplitudes in female mTBI mice compared to female sham mice which suggests a possible potentiation of postsynaptic glutamatergic and GABAergic synaptic function by mTBI in female LHb. However, we found that mTBI resulted in similar shifts in cumulative probability curves of sE/I frequency and synaptic drive ratios to the right without any change in sE/I amplitude curves as seen in male LHb^39^ (Figure 3A-D, Kolmogorov–Smirnov tests, **p<0.01, ****p<0.0001).

**Figure 2.**
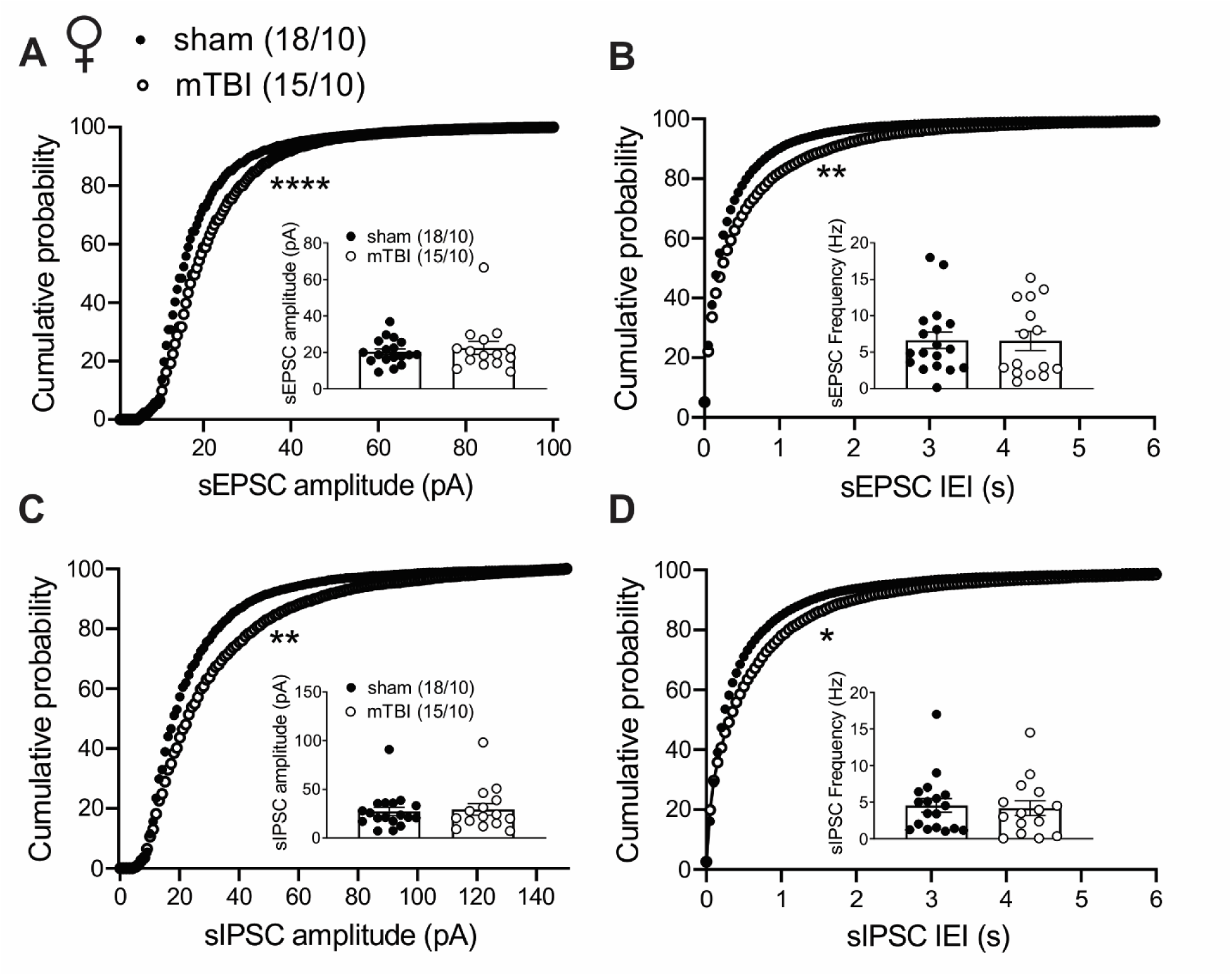
Effects of mTBI on spontaneous synaptic activity in LHb neurons in female mice. A-D show average and cumulative probability amplitude and frequency (inter-event interval) plots of sEPSCs and sIPSCs within the same LHb neurons from female sham (black filled round symbols) and mTBI (black open round symbols) mice at 4 weeks following the injury. mTBI significantly shifted cumulative probability curves of sEPSC and sIPSC amplitude and frequency, resulting in an overall decreased presynaptic spontaneous excitatory and inhibitory transmission in LHb neurons while potentiating postsynaptic spontaneous synaptic activity in female mice. *p<0.05, **p<0.01, ****p<0.0001 by Kolmogorov–Smirnov tests. “(n)” in this and all following graphs represents the number of recorded cells/mice.

**Figure 3.**
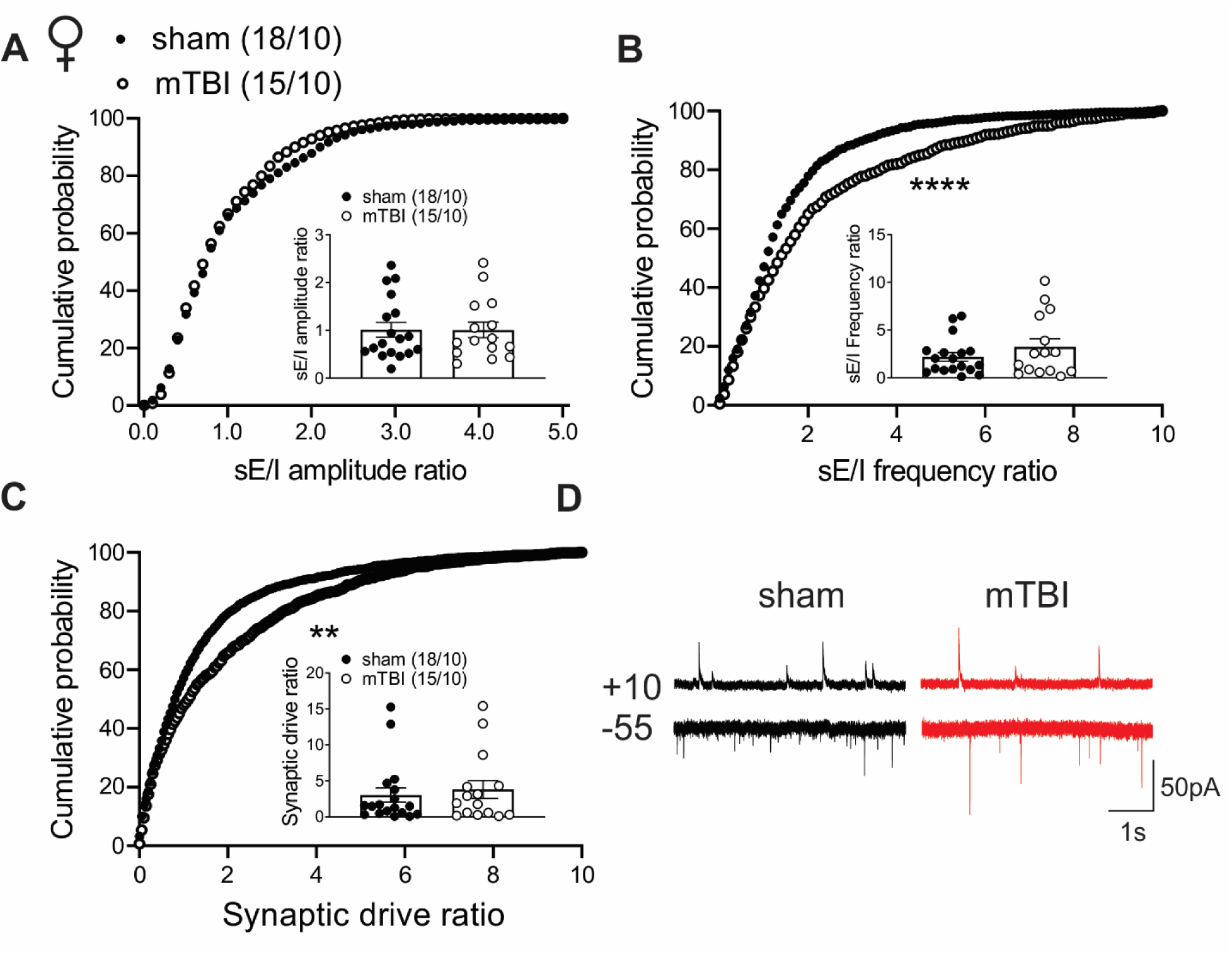
Effects of mTBI on excitation/inhibition balance. **A**, **B**, and **C** show the average and cumulative probability plots of the sE/I amplitude, frequency and excitatory synaptic drive in LHb neurons recorded from female sham (black filled round symbols) and mTBI (black open round symbols) mice. mTBI significantly shifted the distribution curves of sE/I frequency and synaptic drive ratios, resulting in an overall increased excitatory synaptic drive in LHb neurons of female mice. **p<0.01, ****p<0.0001 by Kolmogorov–Smirnov tests. **D** shows representative voltage-clamp recordings of spontaneous excitatory postsynaptic currents (sEPSCs, recorded at –55mV) and spontaneous inhibitory postsynaptic currents (sIPSCs, recorded at +10 mV) within the same LHb neurons in sham (left, black) and mTBI (right, red) mice (calibration bars, 50pA/1 s).

Given that spontaneous synaptic activity recorded as sEPSC and sIPSC are mostly informative of presynaptic component of synaptic transmission including action-potential driven as well as spontaneous neurotransmitter release from presynaptic terminals, our findings highlight that the overall excitatory synaptic drive onto LHb neurons increases following mTBI through a greater suppression of GABA release onto LHb neurons in both male and female mice.

### 3.2. KOR activation within the LHb of sham mice decreases the overall excitatory synaptic drive onto LHb neurons; a neuromodulatory mechanism that is absent following mTBI

To test whether synaptic activity in the mouse LHb can be modulated by Dyn/KOR system, we next examined the effects of bath application of a selective KOR agonist, U50, 488, on spontaneous synaptic transmission onto LHb neurons in LHb-containing slices from male and female sham and mTBI mice. Given the similarity of the general effects of KOR activation on sE/I frequency and excitatory synaptic drive ratios in LHb neurons of male or female mice in either sham or mTBI groups, hereafter we pooled all the KOR-related data from both sexes to generate a larger sample size for statistical power to evaluate differences in KOR stimulation effects between sham and mTBI groups. We found that in general, KOR activation in LHb neurons of non-injured sham mice suppressed only glutamatergic transmission evident from significant reductions in the mean sEPSC amplitude and frequency, a leftward shift in cumulative probability of sEPSC amplitude and a rightward shift in cumulative probability of sEPSC inter-event interval (Figure 4A-D, two tailed paired Student’s t-tests, Kolmogorov-Smirnov tests, *p<0.05, **p<0.01, ****p<0.0001). KOR stimulation in LHb of mTBI mice also resulted in similar reductions in the mean values and shifts in cumulative probabilities of sEPSC amplitude and frequency/inter-event interval suggesting that KOR suppressing effects on presynaptic glutamatergic transmission remained intact in LHb of mTBI mice (Figure 5A-B, two-tailed paired Student’s t-tests, Kolmogorov-Smirnov tests, *p<0.05, ****p<0.0001). However, mTBI additionally reduced the average sIPSC frequency and led to a significant rightward shift in cumulative probability curve of sIPSC inter-event interval in mTBI mice, indicating that mTBI enabled an inhibitory KOR-mediated effect on presynaptic GABA release in LHb neurons of mTBI mice (Figure 5D, two-tailed paired Student’s t-tests, Kolmogorov-Smirnov tests, *p<0.05, ****p<0.0001). Overall, our data indicate that while KOR stimulation presynaptically suppresses only spontaneous glutamatergic transmission onto LHb neurons under normal conditions, mTBI leads to an alteration in KOR modulation of LHb synaptic activity where both presynaptic components of spontaneous glutamatergic and GABAergic transmission onto LHb neurons are sensitive to inhibitory effects of KOR stimulation. Consequently, the overall synaptic effects of KOR stimulation in sham LHb resulted in a significant reduction in the average sE/I frequency and excitatory synaptic drive ratios and leftward shifts in sE/I amplitude, frequency and excitatory synaptic drive (Figure 6A-C, two-tailed paired Student’s t-tests, Kolmogorov-Smirnov tests, *p<0.05, ****p<0.0001). On the other hand, as a result of a dual effects of KOR-mediated suppression of glutamatergic and GABAergic transmission in LHb neurons of mTBI mice, the average and cumulative probabilities of sE/I amplitude, frequency and excitatory synaptic drive onto LHb neurons were unaltered by mTBI (Figure 6D-F).

**Figure 4.**
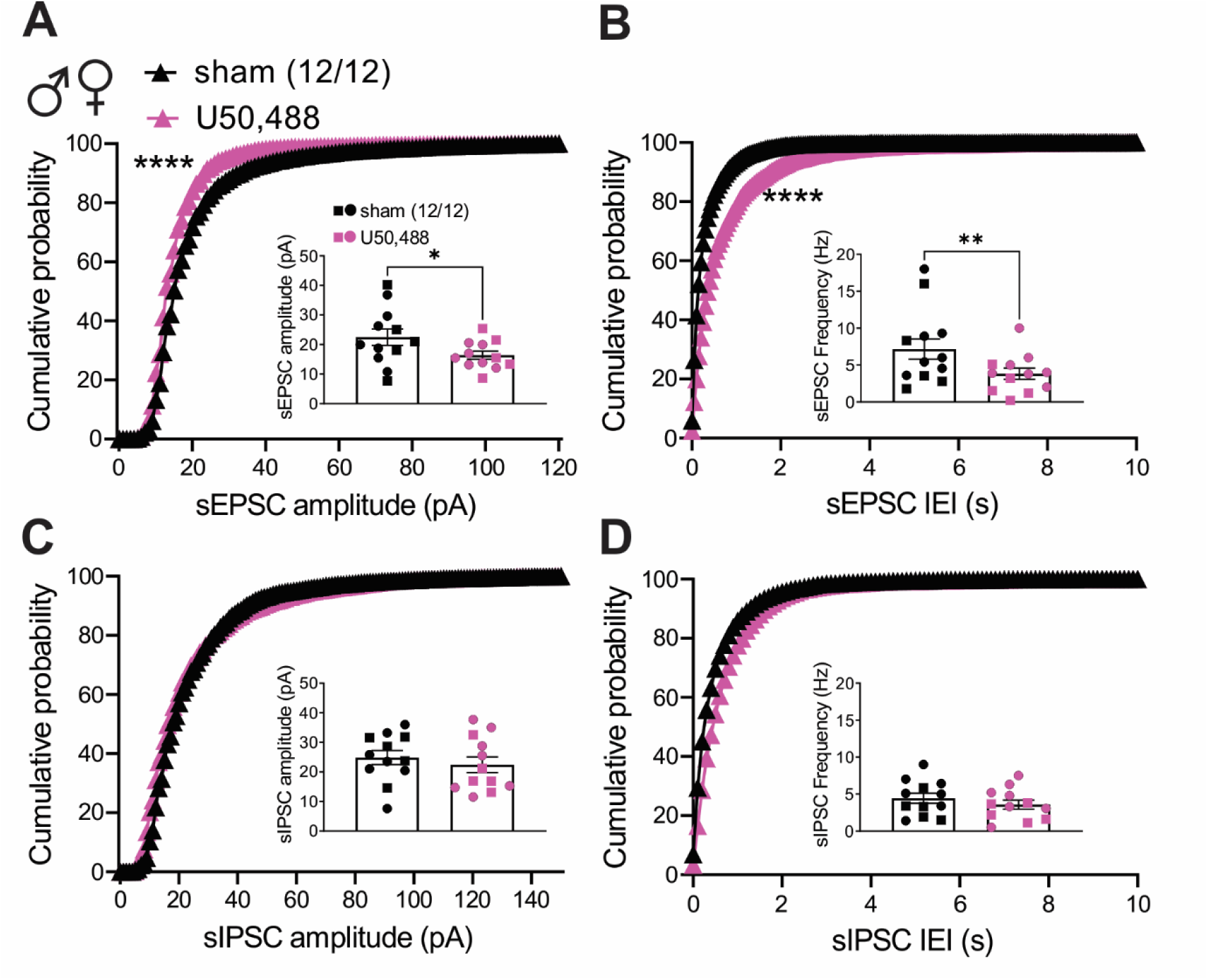
Effects of KOR activation by a selective KOR agonist (U50, 488) on spontaneous synaptic activity in LHb neurons in male and female sham mice. A-D show average and cumulative probability amplitude and frequency (inter-event interval) plots of sEPSCs and sIPSCs within the same LHb neurons from male and female sham mice before (black filled square and round symbols representing data from males and females, respectively) and after bath application of U50,488, 10μM (magenta filled square and round symbols representing data from males and females, respectively) at 4 weeks following the injury. In cumulative probability plots, data from males and females are shown by black filled triangle symbols (before) and magenta filled triangle symbols (after U50,488 application). KOR activation in LHb of sham male and female mice significantly decreased presynaptic and postsynaptic spontaneous excitatory transmission in LHb neurons without any alteration in spontaneous GABAergic transmission. *p<0.05, **p<0.01, ****p<0.0001 by Paired Student t-tests and Kolmogorov–Smirnov tests.

**Figure 5.**
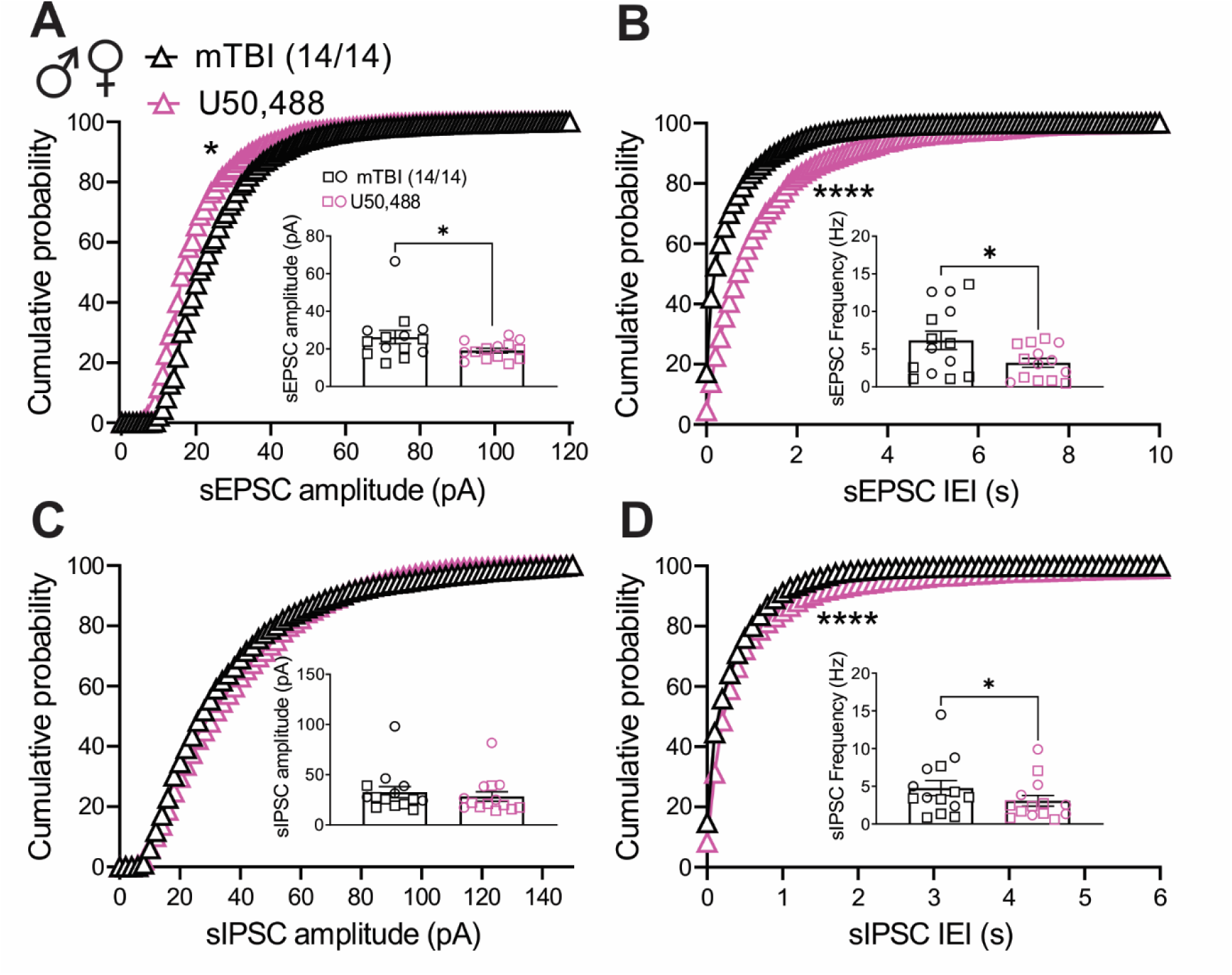
Effects of KOR activation by a selective KOR agonist (U50, 488) on spontaneous synaptic activity in LHb neurons in male and female mTBI mice. A-D show average and cumulative probability amplitude and frequency (inter-event interval) plots of sEPSCs and sIPSCs within the same LHb neurons from male and female mTBI mice before (black open square and round symbols representing data from males and females, respectively) and after bath application of U50,488, 10μM (magenta open square and round symbols representing data from males and females, respectively) at 4 weeks following the injury. In cumulative probability plots, data from males and females are shown by black open triangle symbols (before) and magenta open triangle symbols (after U50,488 application). KOR activation in LHb of mTBI male and female mice significantly decreased presynaptic and postsynaptic spontaneous excitatory transmission and presynaptic GABAergic transmission in LHb neurons. *p<0.05, ****p<0.0001 by Paired Student t-tests and Kolmogorov–Smirnov tests.

**Figure 6.**
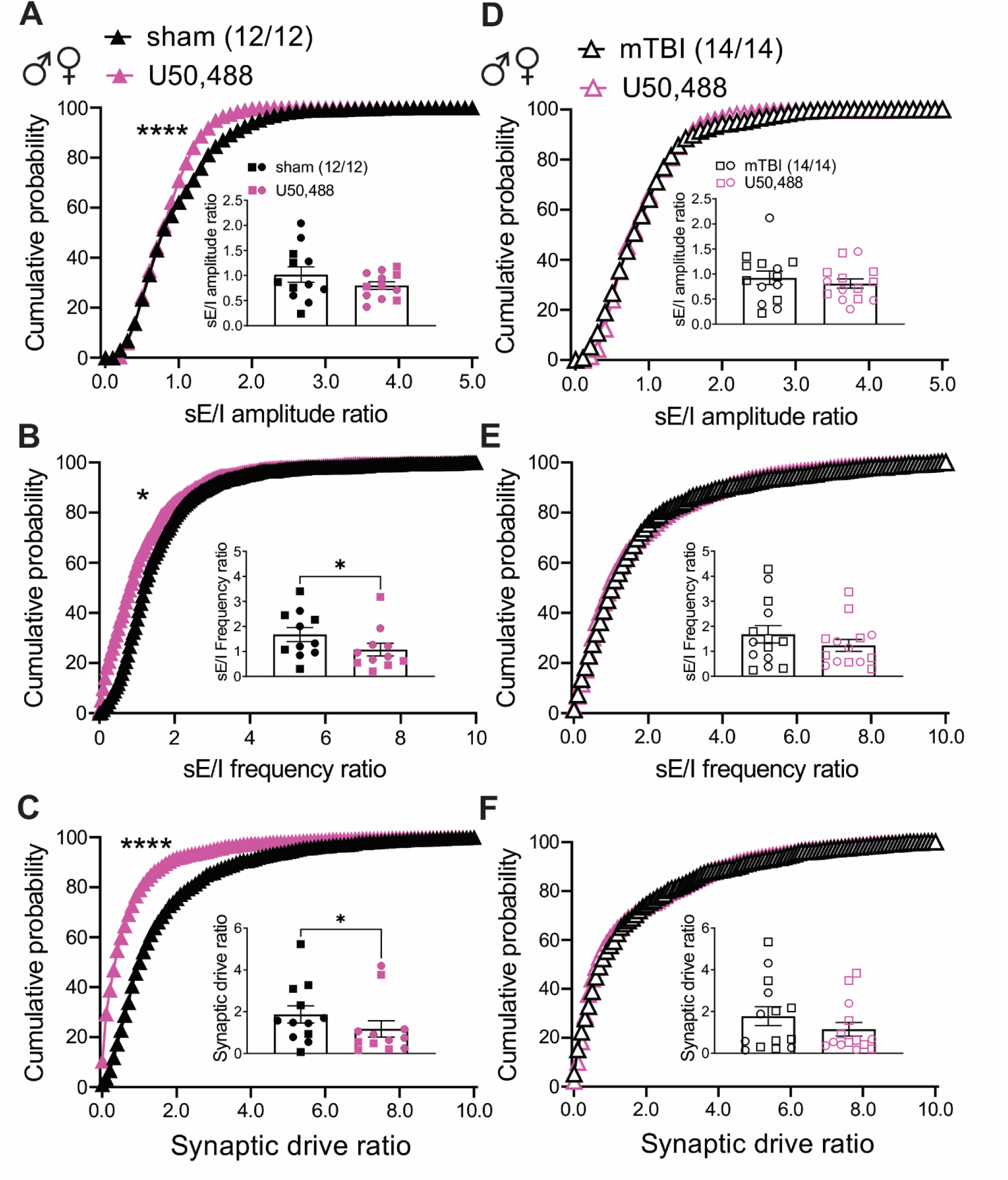
Effects of KOR activation on excitation/inhibition balance in male and female sham and mTBI mice. Figure shows the average and cumulative probability plots of the sE/I amplitude, frequency and excitatory synaptic drive ratios in LHb neurons recorded from male and female sham mice (left panels, **A**-**C**, filled symbols) or male and female mTBI mice (right panel, **D**-**F**, open symbols) before (black) and after U50,488 bath application (magenta). In the average plots square symbols represent data from males, and round symbols show data from females. In cumulative probability plots, data from males and females are shown as triangle symbols. KOR stimulation significantly decreased sE/I amplitude, frequency or excitatory synaptic drive in LHb neurons of male and female sham mice while mTBI mice did not show any alterations in these measurements by KOR stimulation. *p<0.05, ****p<0.0001 by Paired Student’s t-tests and Kolmogorov–Smirnov tests.

Therefore, KOR-mediated neuromodulatory effects in decreasing E/I frequency and synaptic excitatory drive ratios in LHb neurons that shifts the E/I balance towards more synaptic inhibition seems to be lost in mTBI mice.

## 4. Discussion

Here we report the major neuroanatomical sources of Dyn/KOR signaling to LHb and demonstrate that the activation of presynaptic KORs within the LHb provides an inhibitory neuromodulatory braking mechanism mainly by suppression of presynaptic glutamate release, thereby tilting the E/I balance towards inhibition in the mouse LHb. This physiological KOR-mediated inhibitory mechanism could potentially suppress LHb activity or prevent LHb hyperactivity once engaged. We then tested whether the effects of KOR activation on excitatory and inhibitory inputs to LHb neurons were still functional after exposure of male or female mice to repeated closed head injury. We show that mTBI engages presynaptic KORs in a way that KOR activation suppresses not only presynaptic glutamate release but also presynaptic GABA release onto LHb neurons after mTBI. The mTBI-induced additional KOR-mediated inhibition of presynaptic GABA release prevents the KOR-mediated shift in the E/I toward the net inhibition potentially contributing to a mTBI-induced E/I imbalance favoring synaptic excitation and LHb hyperactivity and hyperexcitability that we have observed in male and female mouse LHb^40^.

Our retrograde tracing findings indicate that only mPFC (mPFC^Dyn^) and VMH (VMH^Dyn^) provide direct Dyn projections to the LHb, while diverse cortical and subcortical brain regions that are well-known to provide either glutamatergic and/or GABAergic and/or glutamate/GABA co-releasing synaptic inputs to the LHb express presynaptic KORs in their terminals. The LHb-projecting KOR terminals originated from major projections to the LHb including mPFC, VP, BNST, EP, extended amygdala, different nuclei of hypothalamus, PAG and VTA in addition to the PVT, ZI and midline nuclei (RLi and IF) that we also identified to be KOR-expressing LHb projections. Overall, our anatomical tracing study suggests the possibility of both direct and indirect modulation of LHb through LHb Dyn/KOR systems, either via direct Dyn release from mPFC and VMH (which both also express presynaptic KORs) or through the action of Dyn released within the LHb-projecting brain regions that express presynaptic KORs.

Previously, we found that mTBI increased spontaneous LHb tonic activity via a shift in synaptic E/I balance towards excitation in male mice^39^. Recently, we also found that mTBI induced similar changes in LHb activity and neuronal excitability in male and female mice with increased spontaneous LHb tonic activity in female mice as well as increased LHb excitability in both male and female mice, partly through potentiation of corticotropin releasing factor (CRF)/CRF receptor 1 signaling within the LHb^40^. Here, we confirmed that mTBI similarly resulted in a uniform suppression of spontaneous presynaptic release of glutamate and GABA within the LHb in female mice with a greater suppression of GABA release onto LHb neurons that led to an overall increased excitatory synaptic drive to LHb neurons as we previously reported in male mTBI mice^39, 42^. Interestingly, mTBI also induced a possible compensatory potentiation of both postsynaptic glutamatergic and GABAergic synaptic function in female LHb that was not detected in our previous study in male mice. This may also explain the sex differences in mTBI-induced LHb hyperexcitability where we found male mice exhibited more exaggerated levels of LHb hyperexcitability than females^40^. Nevertheless, the presynaptic effects on spontaneous synaptic transmission following mTBI in females override these postsynaptic compensatory responses, resulting in a bias in E/I balance toward excitation in the female LHb as seen in the male LHb^39^.

Our recent observations of CRF/CRFR1 modulation of neuronal and synaptic activity in the mouse LHb^43^ and mTBI enhancement of CRF/CRFR1 signaling within the LHb which contributes to mTBI-induced LHb hyperexcitability in both male and female mice^40^ suggest that stress neuromodulatory systems including CRF/CRFR 1^43, 44^ and Dyn/KOR signaling^37^ regulating LHb neuronal and synaptic functions may partly contribute to persistent LHb hyperactivity following mTBI. In addition to the extra hypothalamic CRF/CRFR1 neuromodulation of LHb function^40, 43, 44^, we also demonstrated that Dyn/KOR signaling can regulate synaptic and neuronal function in the male rat LHb^37^ although it was unclear if a functional Dyn/KOR system also exist in the mouse LHb and regulate LHb function. Given that Dyn/KOR interactions with CRF signaling within mood– and motivation-related neuronal circuits are shown to mediate behavioral dysregulation of stress and threat responses relevant to neuropsychiatric illnesses^36^, here we then sought to explore the effects of KOR stimulation on spontaneous synaptic activity in LHb neurons in male and female mice and determine how mTBI may alter LHb responsivity to Dyn/KOR system. We show that activation of KORs by a selective KOR agonist (U50,488) only suppressed presynaptic glutamate release without any alterations in spontaneous GABAegic transmission, thereby resulting in a significant decrease in excitatory synaptic drive and regulating synaptic integration by favoring more GABAergic inhibition of LHb neurons in male and female mice. More importantly, we report that mTBI results in a persistent dysregulation of this acute KOR modulation of LHb synaptic integration in which mTBI led to engagement of KOR signaling to additionally suppress presynaptic GABA release in the mouse LHb. Here, we favor the idea that KOR-mediated inhibitory actions on presynaptic glutamatergic transmission within the LHb provides a braking mechanism by which LHb hyperactivity could be prevented and mTBI-loss of this modulatory mechanism could work in parallel with hyperactive LHb CRF/CRFR1 signaling to promote LHb hyperactivity by mTBI. There is some evidence for the possibility of involvement of Dyn/KOR system in long-term aversive and anxiety-like behaviors associated with a repeated blast model of mTBI in male mice where systemic pre-treatment of blast-injured male mice with nor-BNI is shown to prevent behavioral deficits one month post-mbTBI ^38^. Given nor-BNI’s unusual long duration of action where its KOR antagonism can be delayed by hours and persist for months^45^, mTBI-induced mood dysregulation associated with this model potentially requires prolonged KOR antagonism but also our findings highlights the possibility that KOR antagonism in LHb Dyn/KOR system may aggravate the negative impact of mTBI on LHb function and LHb-related mood and motivation regulation while in other mood-related circuits, KOR antagonism may exert antidepressant effects through their Dyn/KOR system. Consistently, activation of KORs is shown to induce dysphoria, negative affective states (depressive and anxiety phenotypes), conditioned place aversion and social avoidance in rodents and/ humans ^46–51^ but preclinical evidence also suggests conflicting anxiogenic and anxiolytic effects for KOR agonists while KOR antagonists reliably exert consistent anxiolytic and antidepressant effects in different animal models ^52^. This contradictory results from KOR activation may be explained by brain-region– and circuit-specific Dyn/KOR modulation of neural circuits regulating mood and motivation including the LHb. Therefore, one possible mechanism for impaired KOR-inhibition of LHb neurons by mTBI could be related to dysregulation of Dyn/KOR modulation of LHb-projecting neurons that co-release glutamate and GABA such as the VTA, EP, LPO, all of which we also found to express KORs through our retrograde tracing. Therefore, mTBI may alter the action of an endogenously released Dyn on the ratio of KOR-mediated inhibition of glutamate versus GABA from these synaptic inputs to the LHb, thereby resulting in synaptic E/I imbalance within LHb.

Interestingly, activation of muscarinic M2 receptors (M2R) is reported to inhibit both presynaptic excitatory and inhibitory inputs to LHb neurons in male and female rats, but with greater inhibition of presynaptic glutamate release resulting in a biased shift of E/I balance toward net inhibition^53^. This M2R-mediated effect on synaptic integration resembles the inhibitory impact of KOR activation on E/I in the mouse LHb. Recent studies from Lupica laboratory have also established the involvement of LHb in suppression of impulsive behaviors through engagement of LHb M2R signaling system ^30, 53^ where they show that impulsivity and impulsive cocaine seeking behaviors are promoted by LHb M2R blockade, possibly through a loss of M2R-mediated synaptic inhibition within the LHb, thereby promoting disinhibition of LHb neurons. Therefore, it will be worthwhile to also consider the possibility that dysregulation of LHb M2 signaling and/or Dyn/KOR plays a role in LHb hyperactivity and behavioral impulsivity following mTBI.

Of high interest for future Dyn/KOR-related circuit-based mTBI studies is the mPFC-LHb pathway considering that the major impact in our mTBI model is delivered to mPFC and our current findings of LHb-projecting mPFC^Dyn^ neurons that may also express KORs. A recent work from Tejada laboratory also demonstrated how pathway-specific Dyn/KOR-mediated modulation of selective synaptic inputs to mPFC and within mPFC microcircuits can regulate E/I balance in mPFC networks and control which limbic inputs can excite mPFC neurons^20^. Most recently, they also elegantly showed the existence of both excitatory and inhibitory Dyn-expressing ventromedial PFC (vmPFC^Dyn^) neurons that can be activated in response to aversive threats (footshocks and footshock-predictive cues) to release Dyn locally within mPFC circuits and limiting passive defensive behaviors (such as freezing)^54^. Although this study did not investigate the long-range projections of the excitatory vmPFC^Dyn^ neurons, given the existence of direct excitatory projections from mPFC to the LHb^55–61^, an established role of LHb in threat-evoked defensive behaviors^62–64^, and our current findings with retrograde tracing studies and mTBI-induced impairment of threat-evoked defensive behaviors, it is plausible to assume that a subset of vmPFC^Dyn^ neurons may also innervate the LHb and provide Dyn release within mPFC-LHb circuits.

## 5. Conclusion

Overall, our study provides evidence for Dyn/KOR-mediated regulation of synaptic integration within the mouse LHb that is susceptible to mTBI injury and may contribute to E/I imbalance and disinhibition of LHb neurons in mice following mTBI. Our tracing and functional studies outlined here provide a framework for future circuit-based studies to determine how LHb Dyn/KOR system acts through distinct LHb circuits that are sensitive or insensitive to KOR activation and control LHb output pathways and LHb-related behavioral responses under normal and pathological conditions.

## Acknowledgments

The opinions and assertions contained herein are the private opinions of the authors and are not to be construed as official or reflecting the views of the Uniformed Services University of the Health Sciences or the Department of Defense or the Government of the United States.

## Authorship contribution

FN, WF, SG and BC were responsible for the study concept and design. WF, SG, SS, FN and ET contributed to the acquisition of animal data. WF, SS, SG, ET, BC and FN assisted with data analysis and interpretation of findings. WF, SG and FN wrote the initial draft of the manuscript. All authors critically reviewed the content and approved final version of manuscript for submission. The authors acknowledge Dr. Yeonho Kim, Dr. Amanda Fu and Laura Tucker at the USU Preclinical Modeling and Behavior Core for supporting the studies.

## Data Sharing

The data that support the findings of this study are available on request from the corresponding author.

